# Development of Equine Polyclonal Antibodies as a Broad-Spectrum Therapy Against SARS-CoV-2 Variants

**DOI:** 10.1101/2022.07.27.501719

**Authors:** Shumin Liao, Yunjiao He, Jing Qu, Yue Shi, Yingzi Liu, Keli Zhao, Junhui Chen, Yue Jing, Clifton Kwang-Fu Shen, Chong Ji, Guxun Luo, Xusheng Zhao, Shuo Li, Yunping Fan, Ziquan Lv, Shisong Fang, Yaqing He, Chunli Wu, Renli Zhang, Xuan Zou, Peng Wang, Liang Li

## Abstract

The Coronavirus disease 19 (COVID-19) pandemic has accumulated over 550 million confirmed cases and more than 6.34 million deaths worldwide. Although vaccinations has largely protected the population through the last two years, the effect of vaccination has been increasingly challenged by the emerging SARS-CoV-2 variants. Although several therapeutics including both monoclonal antibodies and small molecule drugs have been used clinically, high cost, viral escape mutations, and potential side effects have reduced their efficacy. There is an urgent need to develop a low cost treatment with wide-spectrum effect against the novel variants of SARS-CoV-2.

Here we report a product of equine polyclonal antibodies that showed potential broad spectrum neutralization effect against the major variants of SARS-CoV-2. The equine polyclonal antibodies were generated by horse immunization with the receptor binding domain (RBD) of SARS-CoV-2 spike protein and purified from equine serum. A high binding affinity between the generated equine antibodies and the RBD was observed. Although designed against the RBD of the early wild type strain sequenced in 2020, the equine antibodies also showed a highly efficient neutralization capacity against the major variants of SARS-CoV-2, including the recent BA.2 Omicron variant (IC50 =1.867μg/ml) in viral neutralization assay in Vero E6 cells using live virus cultured. The broad-spectrum neutralization capacity of the equine antibodies was further confirmed using pseudovirus neutralization assay covering the major SARS-CoV-2 variants including wild type, alpha, beta, delta, and omicron, showing effective neutralization against all the tested strains. *Ex vivo* reconstructed human respiratory organoids representing nasal, bronchial, and lung epitheliums were employed to test the treatment efficacy of the equine antibodies. Antibody treatment protected the human nasal, bronchial, and lung epithelial organoids against infection of the novel SARS-CoV-2 variants challenging public health, the Delta and Omicron BA.2 isolates, by reducing >95% of the viral load. The equine antibodies were further tested for potential side effects in a mouse model by inhalation and no significant pathological feature was observed.

Equine antibodies, as a mature medical product, have been widely applied in the treatment of infectious diseases for more than a century, which limits the potential side effects and are capable of large scale production at a low cost. A cost-effective, wide-spectrum equine antibody therapy effective against the major SARS-CoV-2 variants can contribute as an affordable therapy to cover a large portion of the world population, and thus potentially reduce the transmission and mutation of SARS-CoV-2.

## Introduction

On 24 November 2021, a novel variant strain of the severe acute respiratory syndrome coronavirus 2 (SARS-CoV-2), B.1.1.529, was first reported as Omicron to the World Health Organization (WHO) by South Africa(1). Soon after two days, the WHO classified B.1.1.529 as a variant of concern (VOC) and designated it as Omicron(2). Sub-lineages of B.1.1.529 including BA.1 and BA.2 emerged and spread quickly around the world, with higher transmissibility and infectivity(3). BA.1 ousted the Delta variant to be the dominating variant of COVID-19 and was replaced by BA.2 before long. In late April 2022, BA.2 was the most dominant variant worldwide(4). Strikingly, new omicron variants are continuously emerging globally. The recently appeared BA.4 and BA.5 variants display higher transmissibility and neutralizing antibody evasion capability, out-competing BA.1 and BA.2(5-7). BA.4/5 subsequently initiated the fifth wave of COVID-19 in South Africa and have been detected in more and more countries worldwide(8).

In these Omicron variants, there are more than 50 amino acid mutations, deletions or insertions compared with ancestral virus strains, especially at the receptor binding motif in the spike protein receptor binding domain (RBD), which exerted concerns of potentially increased transmissibility, reduction in neutralization of spike protein by sera from vaccinated or convalescent individuals, and reduced susceptibility to existing antibody treatments(2, 7). Structural analysis consistently showed that mutations of Omicron variants resulted in reduced affinity between the spike RBD and neutralizing antibodies(3), though the pathogenicity of these Omicron variants was less than early SARS-CoV-2 strains(6). High transmissibility and breakthrough infection of Omicron variants are challenging the efficacy of current therapeutics, including inactivated vaccines like BNT162b2 or mRNA-1273(9-12) and monoclonal antibodies like cilgavimab(8).

Based on the scale of the pandemic, it was estimated that single-point mutations in the large SARS-CoV-2 genome would be generated every day. The pandemic situation calls urgently for effective, specific, and quickly accessible drugs. Polyclonal antibodies from convalescent individuals are commonly used as emergency treatments for emerging infectious diseases. However, the restricted availability and the risk of bloodborne diseases have also typically impeded the widespread clinical applications of the convalescent plasma(13). Equine antibodies, a kind of mature and safe-to-use product with a long history, are cost-effective and affordable in low-income countries where they are needed the most(14-17).

In this study, the RBD of SARS-CoV-2 spike protein was firstly synthesised and purified followed by structural characterization via Circular dichroism spectroscopy and Fourier-transform infrared spectroscopy. We then generated anti-RBD pAbs from equine serum by horse immunization with obtained RBD antigens. The binding affinity of RBD antigens and anti-RBD pAbs were analysed using biolayer interferometry. Several major SARS-CoV-2 variants in the form of both live viruses and pseudoviruses were applied in order to evaluate the spectrum of antibody efficiency. In addition to viral neutralization assay using cell lines, we constructed human respiratory organoids to conduct infection experiments which can better reflect host responses against infection. Three different types of airway organoids, representing nasal, bronchial and lung epithelium, were involved in our study to test the efficacy of the pAbs. Finally, pAbs’ safety was investigated using a mouse inhalation model.

## Materials and Methods

### SARS-CoV-2 spike RBD protein antigen generation

The SARS-CoV-2 spike protein’s receptor binding domain (RBD), Arg319-Phe541 residues, was cloned into a pET21a expression vector (Invitrogen) with a C-terminal 6 His tag. A single colony of the construct was grown in Luria broth (LB) media for protein expression after being converted into bacterial BL21 (DE3)-pLysS competent cells. A high-pressure homogenizer was used to lyse the bacterial pellet. The target protein, containing in inclusion bodies, were washed with urea buffer (2 M) followed by solubilizing with 8M urea-containing buffer (50 mM Tris, pH 9.0, 8M urea, 10 mM beta-ME). Ni^2+^ affinity chromatography and size exclusion chromatography were used to purify denatured protein operating at denaturing conditions. Refolding buffer (50 mM Tris, pH 9.0, 0.4 M arginine, 5 mM GSH, 0.5 mM GSSG) was then prepared to perform protein refolding via fast dilution to decrease the concentration of the urea.

### Circular dichroism spectroscopy

The secondary structure of the generated RBD was measured by the far-UV circular dichroism (CD) spectroscopy, using a Chirascan spectropolarimeter. Refolded RBD was dialyzed into PB buffer (pH 7.4) followed by diluting to a concentration of 0.2 mg/ml for CD measurement. The spectrum was recorded between 260 nm and 190 nm at 20 °C. The spectra represented an average of three individual scans and were corrected for absorbance caused by the buffer. A quartz cuvette with a 0.1 cm path length was used for the measurement. The data was processed and smoothed via the Graphpad Prism 8.0.1 software.

### Fourier-transform infrared spectroscopy

The Fourier-transform infrared (FTIR) spectrometer (Bruker Vertex 70v) coupled with a diamond ATR accessory was used to measure the infrared spectra of RBD using the attenuated total reflectance (ATR) method. Atmospheric effects were eliminated by collecting spectrum without a sample prior to the test. 10 mg/ml RBD protein in PBS buffer or PBS buffer alone were positioned onto the diamond crystal surface. After drying under a stream of nitrogen, the deposited film served as the sample and background respectively. Every spectrum is typically recorded with 256 scans at a resolution of 2 cm^-1^.

### Immunization of horses with RBD

Anti-RBD antibodies were generated by immunizing healthy horses with no detectable antibodies against SARS-CoV-2. The obtained RBD antigens were sterilized by ultra-filtration. Horses were immunized and collected for antiserums. The horse sera before immunization were also collected for negative controls of antibody evaluation. The tilters of serums were tested by an ELISA assay using cPass™ SARS-CoV-2 Neutralization Antibody Detection Kit following standard procedure(18).

### Biolayer interferometry (BLI) analysis of RBD and antibody binding affinity

Anti-RBD-pAbs binding affinities towards the RBD antigen was measured by BLI on an Octet Red 96 instrument (ForteBio, USA) using Ni-NTA biosensors at 30°C with shaking at 1000 rpm. A 60s baseline measurement in kinetics buffer (PBS, 0.02% Tween 20) was conducted firstly followed by loading of RBD antigens (500 nM) for 180 s. Then a second baseline measurement was performed in kinetics buffer for 180 s. 180 s association and 300 s dissociation of anti-RBD pAbs were conducted in different pAbs solutions and kinetics buffer respectively. Results were analyzed by ForteBio Data Analysis software (10.0.3.1). The *K*_*D*_ value of RBD or Tn-RBD binding affinity to pAb was calculated from the binding curves based on the global fit to a 1:1 Langmuir binding model with an R^2^ value of ≥ 0.95. The kinetically derived affinities were calculated as *K*_*D*_ = *K*_*dis*_/*K*_*on*_.

### Cells and viruses

Vero E6 cells (African green monkey kidney cell, ATCC CCL-81) were cultured in complete culture media (Dulbecco’s Modified Eagle Medium (DMEM; Thermo Fisher Scientific, USA) supplemented with 10% fetal bovine serum (FBS) and penicillin (100 U/ml)–streptomycin (100 mg/ml)) at 37°C under 5% CO_2_. SARS-CoV-2/shenzhen/02/2020 (WT), SARS-CoV-2/shenzhen/09/2022(Delta), SARS-CoV-2/shenzhen/08/2022(Omicron BA.1) and SARS-CoV-2/shenzhen/13/2022(Omicron BA.2) were isolated from throat swabs from patients tested positive for SARS-CoV-2. All strains were confirmed by sequencing. Standard plaque assay on Vero E6 cells were conducted for determining virus titers and the virus stocks were stored in aliquots at –80°C until required.

### Neutralization assay

#### Virus

20,000 Vero E6 cells were seeded in each well of a 96-well plate with 100 µl culture media and cultured at 37°C and 5% CO_2_. After overnight culture, cells were then treated with 100 µl virus/antibody mixture solution and incubated for another 1 h. Virus solution was prepared by diluting in culture media allowing the multiplicity of infection (MOI) reach 0.01. Virus/antibody mixture solution was obtained by mixing 60 µl anti -RBD-pAbs solution at serial-diluted concentrations with culture media and 60 µl virus solution. The mixture was incubated at 37°C and 5% CO_2_ for 1h prior to cell infection. The infection media was discarded after infection and the cells were washed with 200 µl PBS twice followed by incubating with carboxymethyl cellulose containing DMEM culture media (#C4888, Sigma). SARS-CoV-2 NP protein in the infected cells was stained for standard ELISpot assay and analyzed by CTL Immunospot analyzer. Infection solution containing pure virus solution (i.e. no antibody) or pure culture media solution were considered virus control and blank control, respectively. Neutralization ratio was determined as:

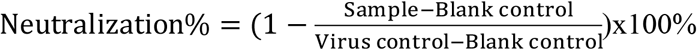

#### Pseudovirus

The pseudovirus including SARS-CoV-2 WT, Delta, Omicron, Beta and Alpha were purchased from Vazyme. 18,000 293T-ACE2 cells were seeded in each well of a 96-well plate with 100 µL culture media. After overnight culture at 37°C and 5% CO_2_, 293T-ACE2 cells were treated with 100 µl pseudovirus/antibody mixture solution and cultured for another 48 h. Pseudovirus solution was prepared by diluting pseudovirus with culture media to a final concentration of 1.3 × 10^4^ TCID_50_/ml. 60 µl anti-RBD-pAb solution were prepared at serial-diluted concentrations in culture media and then mixed with 60 µl pseudovirus solution. The mixture was incubated at 37°C and 5% CO_2_ for 1h prior to cell infection. Neutralizing activity was determined via bioluminescence detection using a microplate reader following the manufacturer’s protocol (Bio-Lite Luciferase Assay System, Vazyme). Briefly, 96-well plates were equilibrated to room temperature first. The culture media was discarded, followed by gentle washing with 100 µL PBS. 100 µl luciferase substrate solution was then added and the bioluminescence was measured after incubating the plate for 10 min at room temperature.

### Construction of 2-D airway organoids

Airway epithelial progenitor cells for 2-D airway organoid culture were derived from patients’ surplus nasal, bronchial or lung biopsy, respectively. The obtained biopsy samples were rinsed with cold DPBS and minced into small pieces for subsequent Dispase I (Stemcell Technologie, CA) dissolving. The harvested cells were cultured in PneumaCult™-Ex Plus Medium (Stemcell Technologie, CA) and then co-cultured with NIH-3T3 feeder cells. When confluent, the cells were transferred to transwells with 0.4µm pores (Corning transwell 3470) and cultured at 37°C under 5% CO_2_. The cells were cultured on an air-liquid interface (ALI) for differentiation in culture media (PneumaCult ALI media, STEMCELL Technologies, Canada) to allow the progenitor cells to differentiate into a mucociliary airway epithelium. Experimental design was reviewed and approved by the ethics committee of the Seventh Affiliated Hospital of Sun Yat-sen University (KY-2021-075-02). Written informed consents were signed by the involved patients as well.

### SARS-CoV-2 infection of 2-D airway organoids

The infection experiments were performed on day 21 of differentiation. 50 µl media containing 10,000 plaque-forming units (PFUs) SARS-CoV-2 was added into the inserts of the transwell plate and incubated for 1 h. The infected organoids in the inserts were then transferred to a new plate after washing for 3 times with PBS. 500 µl serum -free growth medium (05001, 296 STEMCELL Technologies, CA, USA) was added and incubated. 150 µl PBS was added into the transwell’s apical compartment before samples were collected for analysis. Cells and samples, including the PBS wash, were all collected.

### Anti-RBD pAbs intake *in vivo* model

The animal study was approved by Institutional Animal Care and Use Committee, Shenzhen Institute of Advanced Technology, Chinese Academy of Sciences (SIATIACUC-YYS-LL-A0550). Anti-RBD pAbs or saline control were inhaled by 8 week old C57 mice intranasally. Lung samples were collected on day 3 after inhalation for H&E staining.

### Quantification of Viral Gene Expression by qPCR

RNA was extracted from PBS wash described above using TRIzol™ (Invitrogen™, Thermo Fisher Scientific). The extracted RNAs were reversed transcribed into cDNA using a PrimeScrip RT reagent kit (Takara, Japan) and quantified using Coronavirus 2019-nCoV nucleic acid detection kit (fluorescent PCR method) (BioGerm, Shanghai, China) following manufacture’s protocol.

## Results

The applied methodologies for this study were summarized in Figure 1 schematically. Briefly, recombinant SARS-COV-2 spike RBD protein antigens were expressed in *E.coli* cells and then purified chromatographically. Horses were then immunized with the obtained RBD protein antigen to generate the equine anti-RBD serum. The anti-RBD pAbs were isolated from the serum and further sterilized via filtration. Its efficacy were evaluated against SARS-CoV-2 and several variants in 2-D airway organoid models including nasal, bronchial and lung. All of the organoids were infected with live viruses (SARS-CoV-2 WT, Delta, Omicron BA.1and BA.2) and treated separately with mock control and equine anti-RBD pAbs. The virus content in the topical release from each group were measure by qPCR thereby evaluating the treatment effects. Besides, *in vivo* experiments were also conducted for pAbs safety evaluation (**Figure 1a**).

**Figure 1.**
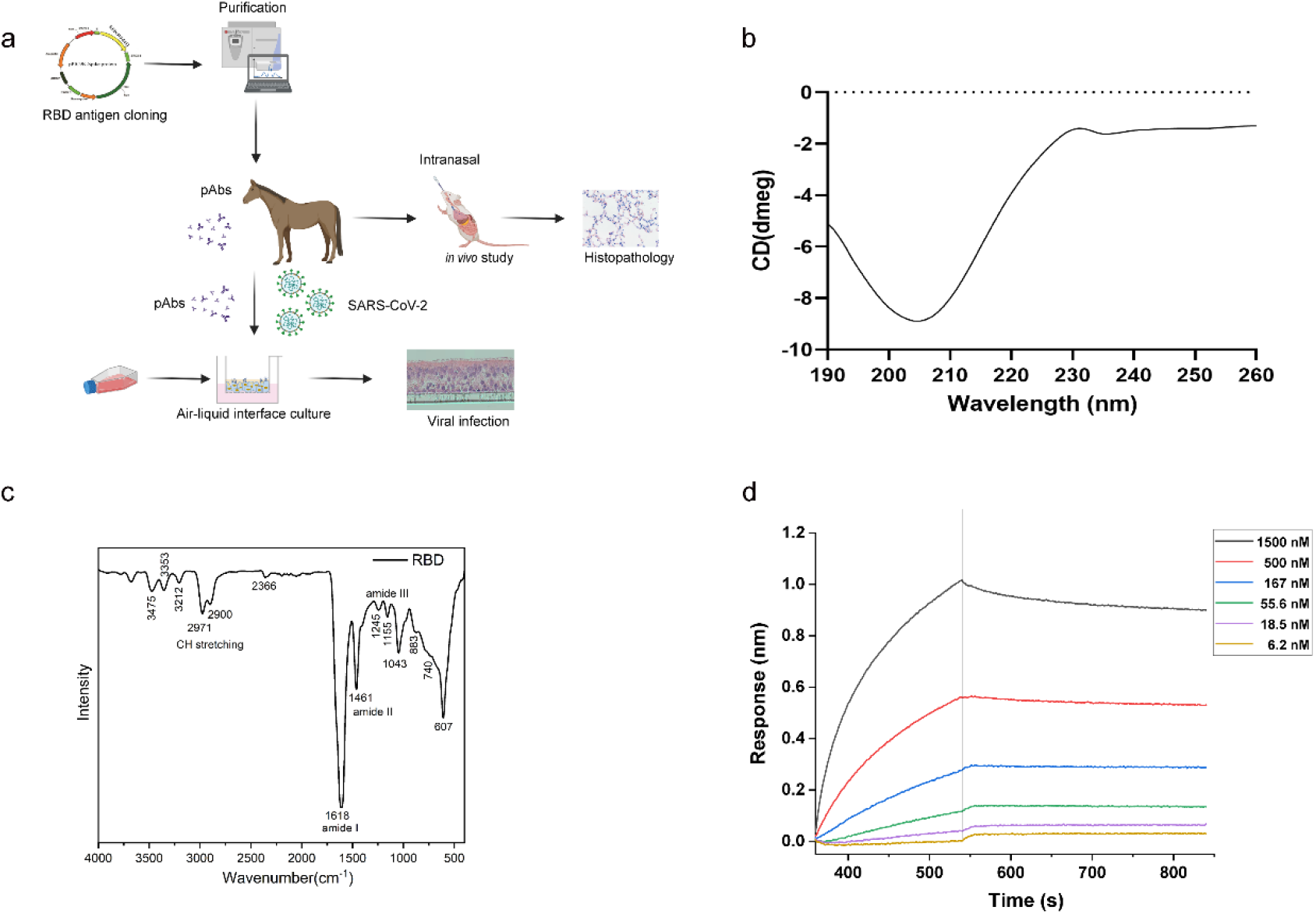
Overview of the study design (a) and RBD characterization (b, c). (a) The RBD antigen was firstly cloned into a vector and purified chromatographically. Then the anti-RBD polyclonal antibodies (pAbs) extracted from horse serum were tested both *in vitro* and *in vivo*. The binding efficacy of antibodies was tested. 2-D airway organoids including nasal, bronchial and lung epitheliums constructed by differentiation in air liquid interphase (ALI) were infected by SARS-CoV-2 and used to evaluate the efficacy of the anti-RBD pAbs.Finally, the anti-RBD pAbs’ safety was tested *in vivo*. The obtained RBD antigen were characterized via CD spectroscopy (b) and FTIR spectroscopy (c) recording the spectrum from 190-260nm and 400-4000nm respectively. (d) The binding curves were generated from basic kinetics analysis of 1500nM, 500nM, 167nM, 55.6nM, 18.5nM and 6.2 nM anti-RBD-pAbs. The association were performed for 550s followed by dissociation in PBS.

### Characterization of receptor-binding domain antigen and binding affinity to pAbs

CD spectroscopy and FTIR spectroscopy were used to structurally characterize the RBD antigens’ structure. As shown in **Figure 1b**, the CD spectrum of *E.coli* produced RBD showed a single minimum at 207 nm, suggesting a characteristic of β-sheet structure. Besides, a maximum around 230 nm was observed, suggesting the contribution of aromatic residues. This is similar to that observed for RBD antigens produced from mammalian or yeast cells(19). The FTIR spectrum (**Figure 1c**) displayed the characteristic peaks at 1614 cm –^1^ (Amide I, C=O vibration), 1467 cm –^1^ (Amide II, N-H stretching) and 2971 cm –^1^ (C-H stretching). Besides, it also showed small peaks around 1100 to 1300 cm–^1^ for Amide III vibration. As compared with the native β-sheet proteins, which typically peak from 1630 to 1643 cm –^1^ for amide I, the shift may be caused by more extended β-sheets or unordered structures.

RBD antigen binding affinity to pAbs were measured by BLI. For all the binding curves, the values increased with the associating time and showed little change curing dissociating process (**Figure 1d**). K_on_ and K_off_, representing the rate constant of association and dissociation respectively, were found the same among all the tested concentrations (1.24E4 Ms^-1^ for K_on_ and 2.56E-4 s^-1^ for K_off_). The results showed that RBD binds to pAb efficiently with a *K*_*D*_ value of 20.6 ± 0.59 nM, indicating a good binding affinity between RBD and equine pAbs.

### Neutralization activity assessment of anti-RBD pAbs

To evaluate the neutralization activity of the anti-RBD pAbs, both virus and pseudovirus were employed in our study. The neutralizing activity of pAbs towards SARS-CoV-2 virus and their mutants were tested using Vero E6 cells and determined by the content of virus nucleocapsid in cells after infection. The IC_50_ values of pAbs towards SARS-CoV-2 WT, Delta, BA.1 and BA.2 mutants were 0.491, 0.254, 0.578 and 1.867 µg/ml, respectively (**Figure 2a**). As to pseudovirus assay, the neutralization activity was assessed using 293T-ACE2 cells by measuring the relative luciferase expression after infection. The IC_50_ values of pAbs against SARS-Cov-2 WT, Delta, Omicron, Beta and Alpha were 0.116, 2.640. 1.190, 13.544 and 3.999 µg/ml respectively. The results suggest our pAbs have a broad-spectrum neutralization capacity towards various SARS-CoV-2 variants in both virus and pseudovirus assays.

**Figure 2.**
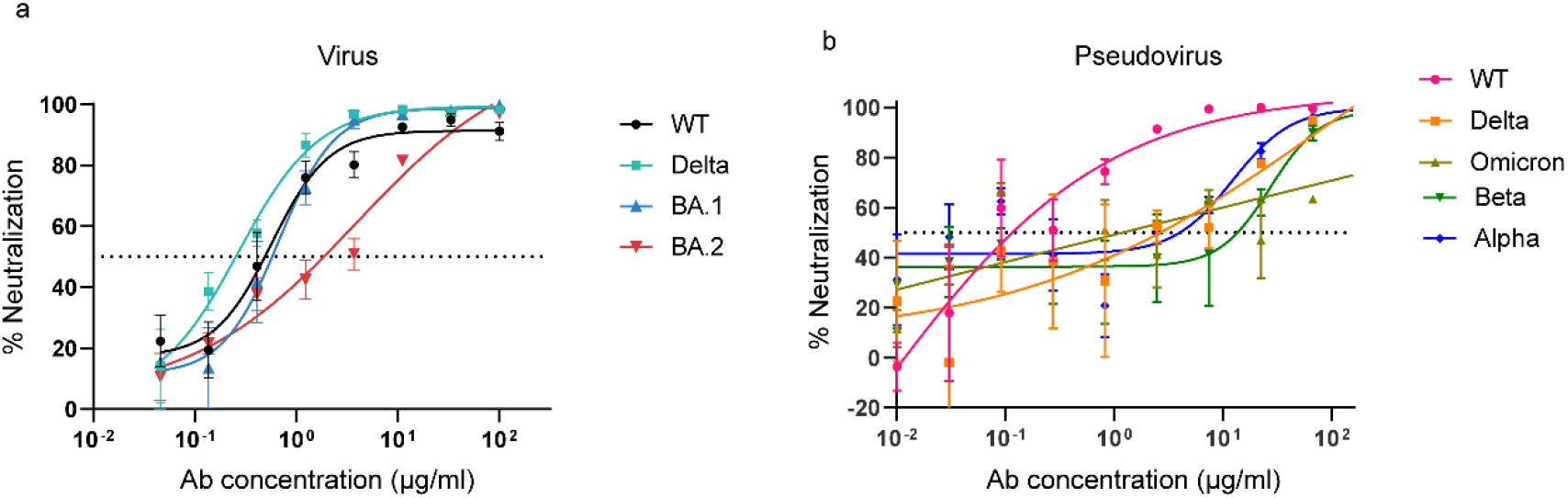
Anti-RBD-pAbs demonstrate potent neutralizing activity towards live virus (SARS-CoV-2 WT, Delta, Omicron BA.1 and BA.2 strains) (a) and pseudovirus (SARS-CoV-2 WT, Delta, Omicron, Beta and Alpha) (b). IC_50_ was determined as the concentration of anti-RBD-pAbs at which 50% of neutralization is reached.

### 2-D airway organoid construction and SARS-CoV-2 infection

Different types of 2-D airway organoids including nasal, bronchial and lung epitheliums were constructed to represent SARS-CoV-2 airway infection. Using nasal organoids as an example, the primary progenitor cells from nasal biopsy were planted in transwell inserts and differentiated in ALI as described in methods section. A pseudostratified epithelium containing goblet cells and ciliated columnar cells were developed and capable of generating mucus. As shown in **Figure 3a**, immunofluorescent staining revealed the apical cells containing both goblet cells (labeled with MUC5AC in red) and ciliated cells (labeled with Ac-α-tubulin in green), indicating the nasal progenitor cells were well differentiated to form a nasal epithelium. Together with the H&E staining (**Figure 3b**) which showed well-organized structures and high similarities compared with nasal biopsy reported in the literatures(20-22), the nasal organoids were confirmed as successfully constructed. The 2-D airway organoids were then applied in the infection model as a testing platform for the neutralization effect of the antibodies. As shown in **Figure 3c** and **3d**, organoids with mock-infection (**Figure 3c**) showed no spike protein expression, while SARS-CoV-2 spike labeled in green were observed in the apical cells (**Figure 3d**) suggested the feasibility of this infection model. We further extended this model for bronchial and lung epitheliums.

**Figure 3.**
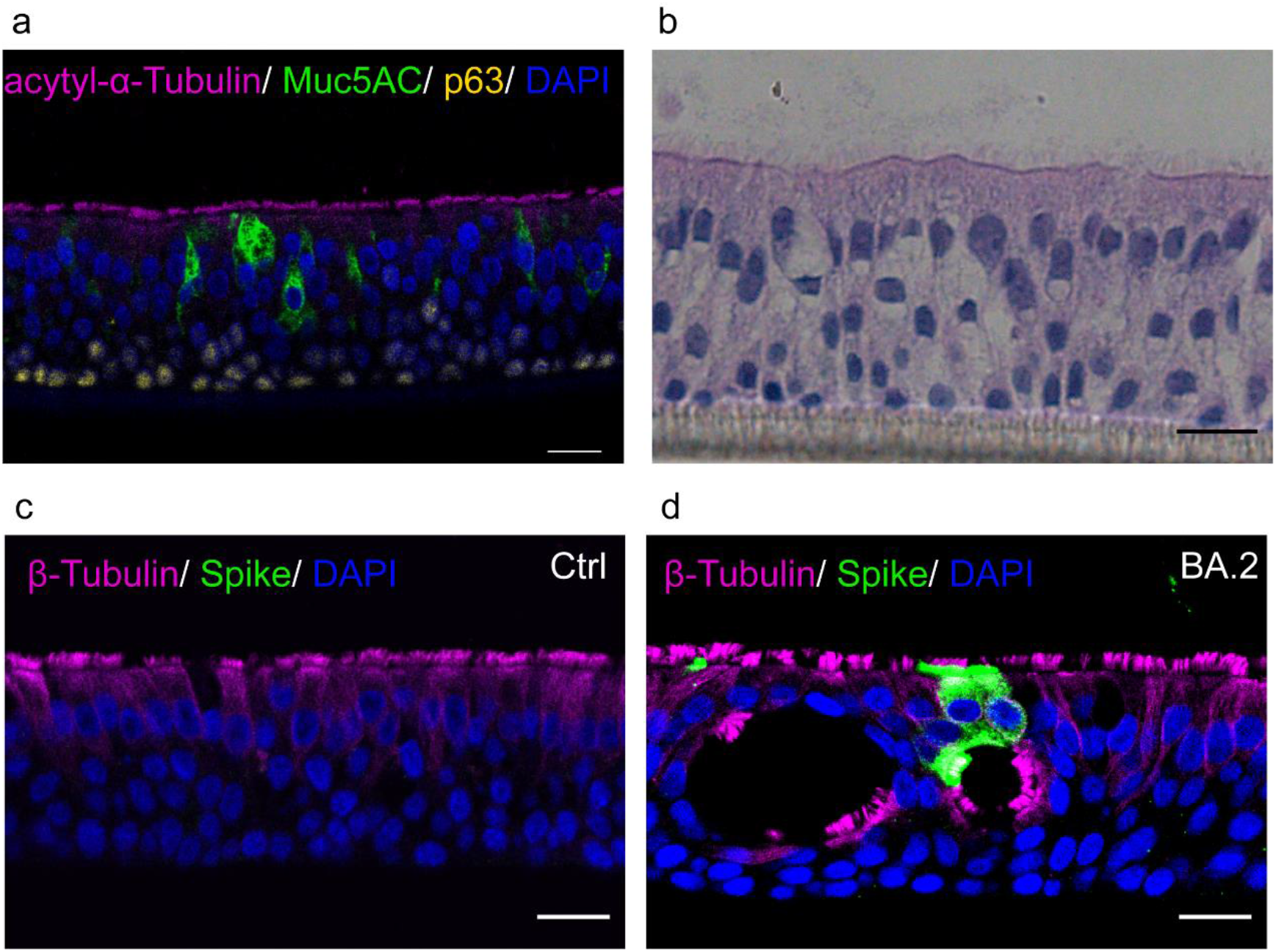
2-D airway organoid and infection model construction from differentiated nasal epitheliums. (a) Representative images of immunofluorescence staining of nasal organoids with ciliated columnar cells stained with AC-α-tubulin (pink), goblet cells stained with Muc5AC (green) and basal cells stained with P63 (yellow). Nucleus were stained by the 4’,6-diamidino-2-phenylidole (DAPI). Scale bar = 20µm. (b) Representative images of hematoxylin-eosin (H&E) staining of nasal organoids. Scale bar = 50 µm. (c, d) Representative immunofluorescence staining of nasal organoids infected with mock control or BA.2. Ciliated columnar cells, virus and nucleus were stained for β-Tubulin (pink), SARS-CoV-2 Spike protein (green) and DAPI (blue) respectively. Scale bar = 20µm.

### Anti-RBD-pAb efficacy and safety assessment

The anti-RBD-pAb efficacy was tested using the organoid infection models constructed. A typical SARS-CoV-2 qPCR kit was used to quantify the topical release of the virus. Delta and BA.2 viral strains were used to infect the organoids constructed to respresent nasal, bronchial and lung epitheliums. As shown in **Figure 4a**, a significant reduction in the ORF1ab and N RNA content were observed in the anti-RBD pAb-treated groups (Inf+Ab) among all organoid models for SARS-CoV-2 infection compared with their non-treated infection counterparts. **Figure 4b** illustrated similar trends for BA.2-infected organoids that the virus content was decreased in pAb-treated groups. We then applied mouse inhalation model to evaluate the safety of our pAbs. As shown in **Figure 4c** and **4d**, H&E staining of lung sections after either saline or anti-RBD pAbs inhalation showed no significant immune cell infiltration in the mouse lungs, demonstrating that the pAbs did not induce significant immune responses in the mouse model.

**Figure 4.**
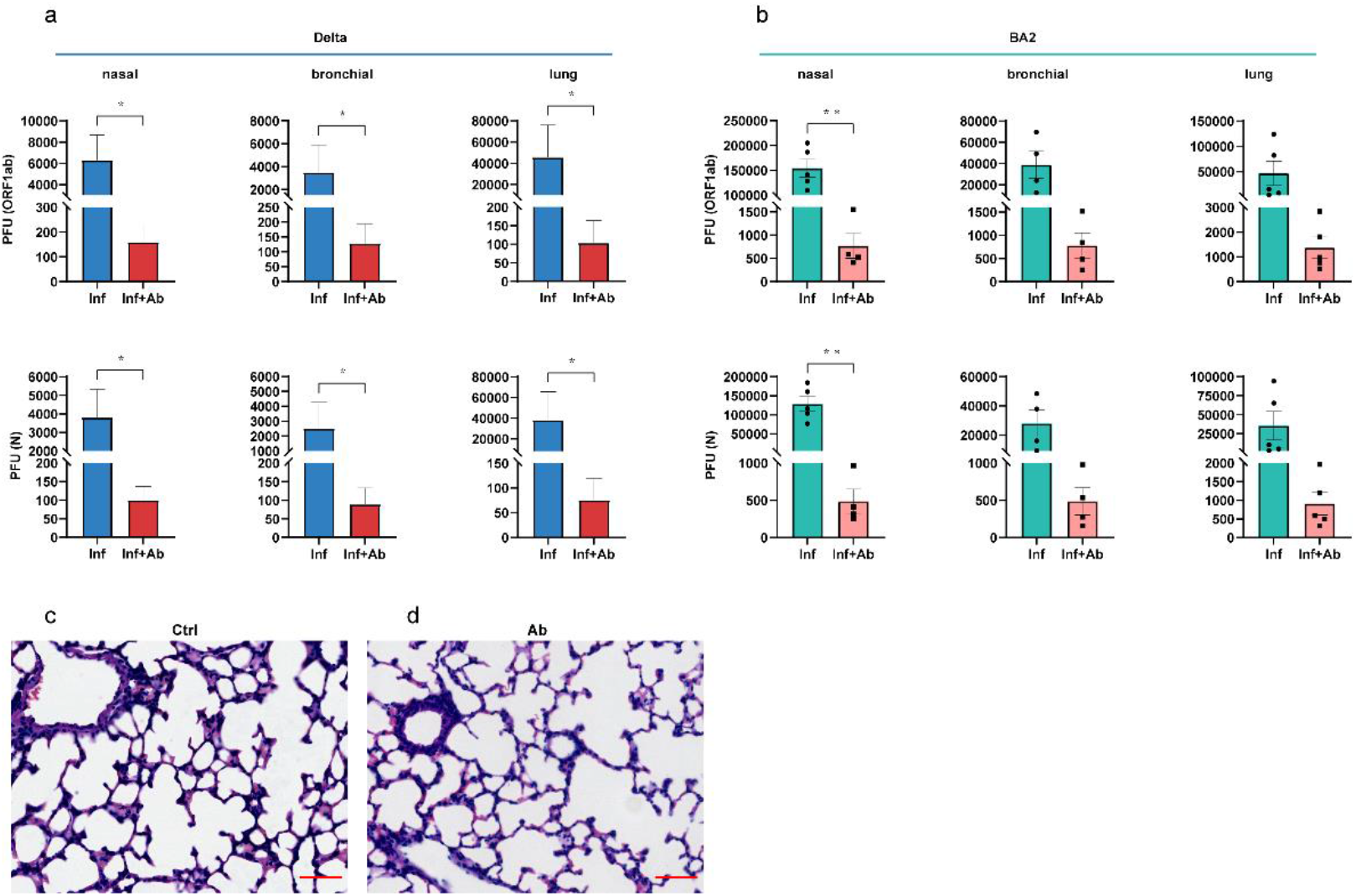
Anti-RBD-pAb efficiency and safety tests. qPCR quantification of ORF1ab and N domains of SARS-CoV-2 Delta (a) and BA.2 (b) strains in topical secretions of the infected organoids as indicated. H&E-stained lung sections from mice with inhalation of saline control (c) or anti-RBD pAbs (d) showed no significant difference. Scale bar = 50µm.

## Discussion

COVID-19 caused by SARS-CoV-2 has emerged worldwide as an unprecedented public health emergency for more than 2 years. Vaccine development has been considered as the best long-term solution to this pandemic. To date, many types of SARS-CoV-2 vaccines have been licensed and administered(23). However, SARS-CoV-2 still poses significant health challenges globally even after boost doses have been required for nearly a year. This is mainly due to the constant emerge of new variants of SARS-CoV-2, the uneven accessibility of vaccines, especially for developing countries, and the effectiveness and side effect concerns of vaccines among the pregnant, the elderly and the immunocompromised populations. This highlights the critical needs for a broad-spectrum treatment capable of large scale production at relatively low cost(24). With decades of experience on equine antibodies production, we conducted SARS-CoV-2 antibody production via horse immunization with the RBD antigens of the SARS-CoV-2 spike protein followed by antibody purification. Our obtained anti-RBD pAbs were then tested against the major variants of SARS-CoV-2 and showed potent neutralization capability against both live virus and pesudovirus (**Figure 2**). Their excellent efficacy were also demonstrated on several respiratory organoid SARS-CoV-2 infection models including nasal, bronchial and lung organoids, showing significant blockage of viral infection and reduction of viral replication upon treatment by equine polyclonal antibodies (**Figure 4**). Our study demonstrated equine anti-RBD pAbs had great potential as a broad-spectrum and cost-effective drug. The manufacture of equine antibody drugs has already been well established, with mature production lines and production processes, making side effects manageable. Besides, the overall development allowing mass production of new equine antibodies within 6 months enables the quick adaption towards new variants when needed, further making them a promising solution for this and future possible pandemics.

In terms of safety and effectiveness for long term usage, equine products had showed their advantages over vaccines and small molecular drugs in several aspects including drug resistance and evasion by novel variants, hepatotoxicity and nephrotoxicity. In late 2021, multiple new variants of SARS-CoV-2, including the alpha, beta, delta and omicron variants, have drawn great concern due to their enhanced transmissibility, virulence and the potential of immune evasion from the host defense built by both vaccinations and previous infections of SARS-CoV-2. More lines of evidence have demonstrated the decreased protection from some vaccine products towards these mutants(25-28). The hindered accessibility to COVID-19 vaccines in low-income countries caused differences in the levels of inoculation, which could also lead to virus mutations and new variants(29). As to small molecule drugs, their liver and kidney toxicity, relatively long development timeline and drug resistance issues were the general concerns raised. Given the past history of small molecular drug development for influenza virus, the continuous evolvement of influenza virus caused rapid emergence of resistance to existing drugs, particularly to adamantanes, followed by oseltamivir, highlighting the constant requirements for new drug development(30). It could be anticipated that the widely disseminated SARS-CoV-2 virus will also develop drug resistance problems within a relatively short time frame due to its wide spread globally. Equine antibodies, as mature antibody therapeutics, can overcome these limitations. Their safety has been recognized for ages by World Health Organization over several widespread diseases(31). In this paper, our results showed the possibilities of using horse immunization to generate anti-RBD pAbs with excellent neutralization activities against several wide spread SARS-CoV-2 variants including Delta, BA.1 and BA.2, on top of WT. Besides, the neutralization effects were also observed using nasal, bronchial and lung epithelial organoid infection models that significant reduction of virus content were shown among all the tested organoids. Our *in vivo* study by a mice inhalation model found no significant differences in lung tissue morphology between a saline control and the antibody treatment (**Figure 4**), confirming on the safety of the anti-RBD pAbs.

Equine antibodies are cost-effective therapeutic products compared to other antibodies. Using polyclonal antibodies derived from horses for the treatment of diseases like rabies, tetanus, diphtheria etc. is a well-known and easily scalable technology owing a history of over a century across the world(32-37). In our study, after horse immunization, large amounts of sera can be collected monthly in a repeated manner to generate high titers of neutralizing antibodies. The harvested antibodies showed high binding affinity towards RBD antigens and potent neutralization activity towards both live virus and pesudovirus of several major variants of SARS-CoV-2. Since equine relevant products are routinely produced in developing countries, it could be easily produced in many parts of the world to ensure the accessibility of effective COVID-19 treatments especially for low-income population.

The proposed administration routine of our equine anti-RBD pAbs includes both inhalation and injection in future clinical applications. The inhalation method exhibit various advantages on prophylactic and therapeutic applications(38). For example, inhalation can block the transmission route among individuals by attaching the antibodies on the respiratory tract surfaces, which can prevent virus invasion into the airway cells and also prevent the virus from being coughed up by the patient. Besides, there are less side effects for the inhalation administration due to its topical medication in the airway. Considering neutralizing antibodies should block the internalization of the virus, early stage and asymptomatic patients could also benefit to prevent severe illness and viral transmission, which is enabled by the low cost and high safety of the equine pAbs.

The possible reason of our equine anti-RBD pAbs having broad spectrum neutralization effect towards the major SARS-CoV-2 variants is potentially due to the intrinsic complex immune responses in horses tending to develop broad spectrum avidity than monoclonal antibodies for their cognate antigens(39). pAbs recognize a vast array of epitopes, reducing the risk of vial escape mutations. In addition, healthy horses generally had better tolerance towards the side effects of immunization by antigens than human beings, especially for the elderly, pregnant and immunocompromised population. We believe with the advances in the vaccine technology development such as nucleic acid vaccines, the production of equine antibodies against possible new mutant strains using equine immunization by new forms of vaccine-like products can be further accelerated. This low-cost, broad-spectrum and highly productive equine antibodies could be beneficial to a broader population including people who respond inefficiently to vaccines and people who cannot afford expensive medications.

## Acknowledgments

The study was supported by the National Natural Science Foundation of China (81900071), the Natural Science Foundation of Guangdong Province of China (2021A1515010004), Shenzhen Science and Technology Program (JCYJ20190809143601759) and SUSTech Starting-up Research Grant (Y01416133).

Figure 1A was created using BioRender.com.

## Author Contributions

Study design: LL, PW, ZX, JQ; Experiments: SML, YJH, JQ, YZL, KLZ, JHC, YJ, CKS, CJ, GXL, SL, YPF, ZQL, SSF, YQH, XSZ, CLW, RLZ; Data analysis: SML, YJH, YS, JQ, LL; Manuscript: SML, YS, LL.

